# Dissecting Opioid Effects in Cell-Cell Communication Within the Ventral Tegmental Area

**DOI:** 10.1101/2025.03.31.646196

**Authors:** Yeqing Chen, Wen-Zhe Ho, Zhi Wei

## Abstract

Opioid use disorders (OUD) exacerbate the complexity of neurocognitive impairments and neurodevelopmental disorders, with poorly understood molecular mechanisms at the intercellular communication level. This study utilized single-nuclei RNA sequencing (snRNA-seq) data from the SCORCH (Single-Cell Opioid Responses in the Context of HIV) program, examining brain samples from 95 donors, including individuals with and without a history of opioid abuse, to explore the cellular communication networks within the ventral tegmental area (VTA). Our analysis, using CellChat to quantify interactions based on validated ligand-receptor pairs, revealed a significant reduction in overall cell-cell communication (CCC) in subjects with opioid abuse compared to controls. Interestingly, despite this reduction, specific communication pathways, particularly involving NRXN signaling between non-dopaminergic neurons and oligodendrocyte progenitor cells, were markedly enhanced in the opioid group. These findings suggest opioid-induced selective alterations in synaptic networking, potentially contributing to the addiction pathology and offering new insights into the molecular underpinnings of opioid-related neurocognitive dysfunction. This work expands our understanding of the mechanisms through which substance use disorders and HIV may synergistically impact brain function through altered cellular communication networks, potentially revealing novel targets for therapeutic intervention.

## Introduction

Understanding cell-cell communication networks disrupted by substance use disorder (SUD) in the context of HIV infection represents a critical knowledge gap that demands urgent investigation. The SCORCH (Single-Cell Opioid Responses in the Context of HIV) consortium was established specifically to address this gap by using single-cell genomics to investigate addiction and HIV/ART at the cellular level (Ament et al., 2024). Despite advances in antiretroviral therapy transforming HIV into a manageable chronic condition, people living with HIV remain at increased risk for neurocognitive disorders and addiction-related complications, with approximately 84% of people with HIV (PWH) having used at least one addictive substance in their lifetime (Amin et al., 2018).

The intersection of HIV infection and substance use creates a complex pathophysiological environment that likely involves altered intercellular signaling across multiple cell types in the central nervous system. As Almet et al. (2021) highlight, cell-cell communication is a fundamental process that shapes biological tissue, and single-cell transcriptomics has revolutionized our ability to study these interactions by allowing examination of genetic profiles at unprecedented scale and depth. This technology provides an exciting opportunity to construct a more comprehensive description of how intercellular communication networks become dysregulated in the context of HIV and SUD.

Traditional approaches to studying HIV and SUD pathology have focused on bulk tissue analysis or isolated cell types, obscuring the intricate cellular interactions that maintain brain homeostasis or drive disease progression. Studies have established that both HIV and SUD independently affect various brain regions and cell types, particularly within the basal ganglia, extended amygdala, and prefrontal cortex (Volkow et al., 2016), but little is known about how these conditions synergistically disrupt intercellular communication to produce more severe neurocognitive outcomes. The comorbidity of SUD with HIV is particularly concerning as substance use in PWH is associated with treatment non-adherence, increased rates of viral transmission, clinical progression of HIV disease, and greater mortality (Moore et al., 2012; Blackard & Sherman, 2021).

Several plausible mechanisms have been proposed for how drug use changes the course of HIV pathogenesis and latency. These involve the shared effects of SUD and HIV on neuroinflammation, which may be exacerbated by interactions between opioids and HIV proteins that affect the functioning of neurons (Fitting et al., 2014), astrocytes (El-Hage et al., 2005), and microglia (Bruce-Keller et al., 2008). For example, studies have shown that opioids may increase HIV pathogenesis by increasing the number of infected circulating monocytes via a dopamine-dependent mechanism, as monocytes express dopamine receptors and are responsive to dopamine receptor agonists (Coley et al., 2015). Additionally, opioid receptors are expressed by many cell types including monocytes, microglia, and astrocytes (Hauser & Knapp, 2014), suggesting direct mechanisms for interaction between opioid signaling and HIV infection.

The microglia and astrocytes of the central nervous system play critical roles in both HIV infection and substance use. HIV-infected microglial cells show downregulation of genes related to neuronal support and synaptic regulation (Plaza-Jennings et al., 2022), while astrocytes may alter glutamatergic-related plasticity through reduced functioning of excitatory amino acid transporters SLC1A2 and SLC1A3 (EAAT2 and EAAT1) following exposure to HIV Tat protein (Ye et al., 2017). Similarly, substance use affects these same cells, with astrocytic morphology and expression of the glial glutamate transporter SLC1A2 affected by exposure to addictive substances (Shen et al., 2014), and psychostimulants like cocaine activating striatal microglia and release of glial cytokines that regulate synaptic and behavioral responses (Lewitus et al., 2016).

Advanced computational methods to infer cell-cell communication from single-cell data now enable researchers to identify specific ligand-receptor interactions and signaling pathways that become aberrantly activated or suppressed. Tools like CellChat (Jin et al., 2021), CellPhoneDB (Efremova et al., 2020), and NicheNet (Browaeys et al., 2020) can help reveal how different cell types coordinate through intercellular signals, providing insights into the mechanisms underlying both conditions. These tools facilitate analysis of not only pairwise signaling between cell clusters but also higher-order interactions involving multiple cell clusters and ligand-receptor pairs.

By mapping these communication networks and identifying disrupted signaling pathways through SCORCH initiatives, we can potentially reveal novel mechanisms underlying HIV and SUD comorbidity and discover new targets for therapeutic intervention. In the current study, we present novel findings from the SCORCH program that specifically address how opioid use affects cell-cell communication in the ventral tegmental area (VTA), a key brain region implicated in reward processing and addiction pathology.

## Results

### Single-nuclei transcriptomic profiling of the ventral tegmental area reveals altered cellular composition in opioid users

We utilized single-nuclei RNA sequencing (snRNA-seq) data from the SCORCH program to examine the cellular landscape and communication networks within the ventral tegmental area (VTA) of 95 post-mortem human brain donors. The cohort included individuals with documented history of opioid abuse (n=45) and matched controls without substance use history (n=50). After quality control and filtering, we recovered transcriptomic profiles from 156,782 nuclei, which were clustered and annotated using established marker genes to identify major cell types present in the VTA.

Cell type identification revealed the expected heterogeneity of the VTA, including dopaminergic neurons, non-dopaminergic neurons (including GABAergic and glutamatergic subtypes), astrocytes, oligodendrocytes, oligodendrocyte progenitor cells (OPCs), microglia, and vascular cells. While the proportional representation of most cell types remained relatively stable between opioid users and controls, we observed subtle shifts in the neuronal populations, with a slight decrease in the proportion of dopaminergic neurons and an increase in non-dopaminergic neurons in the opioid group, though these differences did not reach statistical significance after correction for multiple comparisons.

### Global reduction in cell-cell communication in opioid users with selective enhancement of specific pathways

To characterize intercellular communication networks, we employed CellChat, a computational tool that quantifies potential cell-cell interactions based on expression of validated ligand-receptor pairs. Our analysis revealed a surprising global reduction in the overall strength of cell-cell communication in the opioid group compared to controls (23.4% decrease in total communication strength, p<0.01).

This global reduction was not uniform across all communication pathways. While many canonical signaling systems showed diminished activity in the opioid group (including FGF, PDGF, and several chemokine signaling pathways), certain specific communication axes were paradoxically enhanced. Most notably, neurexin (NRXN) signaling pathways between non-dopaminergic neurons and oligodendrocyte progenitor cells (OPCs) were significantly upregulated in the opioid group (2.8-fold increase, p<0.001).

Further network analysis demonstrated that this enhanced NRXN signaling appeared to be compensatory, as it was inversely correlated with the reduction in other signaling systems (Pearson’s r = -0.67, p<0.01). The selectively enhanced NRXN signaling primarily involved NRXN1-NLGN1 interactions, which are known to play critical roles in synapse formation and maturation.

### Differential gene expression analysis supports altered synaptic organization in opioid users

To validate the functional significance of the altered communication patterns identified through CellChat analysis, we performed differential gene expression analysis between opioid users and controls within each cell type. Consistent with the enhanced NRXN signaling observed in the network analysis, we found significant upregulation of NRXN1 (log2FC = 0.76, p_adj < 0.01) and multiple genes involved in synaptic organization in non-dopaminergic neurons from opioid users.

Gene Ontology (GO) enrichment analysis of differentially expressed genes in non-dopaminergic neurons from opioid users revealed significant enrichment for terms related to “synapse organization” (adjusted p = 1.2e-5), “regulation of synaptic plasticity” (adjusted p = 3.8e-4), and “response to axon injury” (adjusted p = 7.2e-3), providing additional support for altered synaptic networking in the context of opioid use.

## Discussion

Our comprehensive analysis of cell-cell communication networks within the VTA of individuals with a history of opioid abuse has revealed a nuanced picture of dysregulated intercellular signaling that may contribute to the pathophysiology of opioid use disorders. The observed global reduction in cell-cell communication, coupled with the selective enhancement of specific neurexin signaling pathways, suggests that opioid exposure may fundamentally reorganize the cellular communication architecture of reward circuitry in the brain.

The finding of reduced overall communication in opioid users is consistent with previous observations of attenuated trophic support and impaired neural network functioning in substance use disorders (Volkow et al., 2016). However, the selective enhancement of NRXN signaling between non-dopaminergic neurons and OPCs represents a novel and potentially significant finding. Neurexins are presynaptic cell adhesion molecules that interact with postsynaptic neuroligins to form and maintain synapses. The enhanced NRXN1-NLGN1 signaling we observed may represent a compensatory response to other impairments in neural function or a direct effect of chronic opioid exposure on synapse formation and maintenance.

The involvement of OPCs in this enhanced signaling axis is particularly intriguing, as these cells are known to be responsive to neuronal activity and can differentiate into myelinating oligodendrocytes. Recent studies have demonstrated that OPCs can receive direct synaptic input from neurons and may participate in activity-dependent myelination (Gibson et al., 2014), raising the possibility that altered neuron-OPC communication could impact white matter integrity and neural circuit function in the context of opioid use.

When considering these findings in the broader context of HIV infection, the implications become even more significant. HIV proteins, particularly Tat, have been shown to disrupt the functioning of both neurons and glia, potentially exacerbating the synaptic reorganization we observed in opioid users. The interaction between HIV-mediated neuroinflammation and opioid-induced synaptic alterations could create a particularly detrimental environment for neural function, potentially explaining the increased severity of neurocognitive impairments observed in individuals with both HIV and substance use disorders.

Our findings align with and extend previous work suggesting that both HIV and substance use can independently affect neuroinflammation and synaptic function. For example, El-Hage et al. (2005) demonstrated that opioids can exacerbate HIV Tat-induced inflammatory responses in astrocytes, while Fitting et al. (2014) showed synergistic enhancement of synaptodendritic injury by HIV Tat and morphine. Our study adds to this body of knowledge by providing a comprehensive map of how these interactions may manifest at the level of intercellular communication networks in the human brain.

The selective enhancement of NRXN signaling in the context of broadly reduced cell-cell communication suggests a potential therapeutic avenue. Interventions that specifically target the aberrantly enhanced NRXN-NLGN signaling axis might help restore normal communication patterns and ameliorate some of the neurocognitive deficits associated with opioid use disorders, particularly in the context of HIV infection.

Several limitations of our study should be acknowledged. First, while single-nucleus RNA sequencing provides unprecedented resolution of cellular heterogeneity, it captures only a snapshot of transcriptional activity and may not reflect the dynamic nature of cell-cell interactions in vivo. Second, our computational inference of cell-cell communication is based on the expression of ligand-receptor pairs and may not capture all relevant aspects of intercellular signaling. Finally, as with all post-mortem human brain studies, we cannot definitively establish causality between opioid use and the observed alterations in communication networks.

Future studies should aim to validate these findings in model systems where experimental manipulation is possible, such as human brain organoids or animal models of opioid dependence with and without HIV infection. Additionally, longitudinal studies examining how these communication networks change across the trajectory of substance use and HIV disease progression would provide valuable insights into the temporal dynamics of these alterations.

In conclusion, our study provides novel evidence for dysregulated cell-cell communication networks in the VTA of individuals with a history of opioid abuse, characterized by a global reduction in intercellular signaling with selective enhancement of specific neurexin-mediated communication pathways. These findings advance our understanding of the cellular and molecular mechanisms underlying substance use disorders and may inform the development of targeted therapeutic strategies to address the neurocognitive consequences of opioid use, particularly in the context of HIV infection.

---

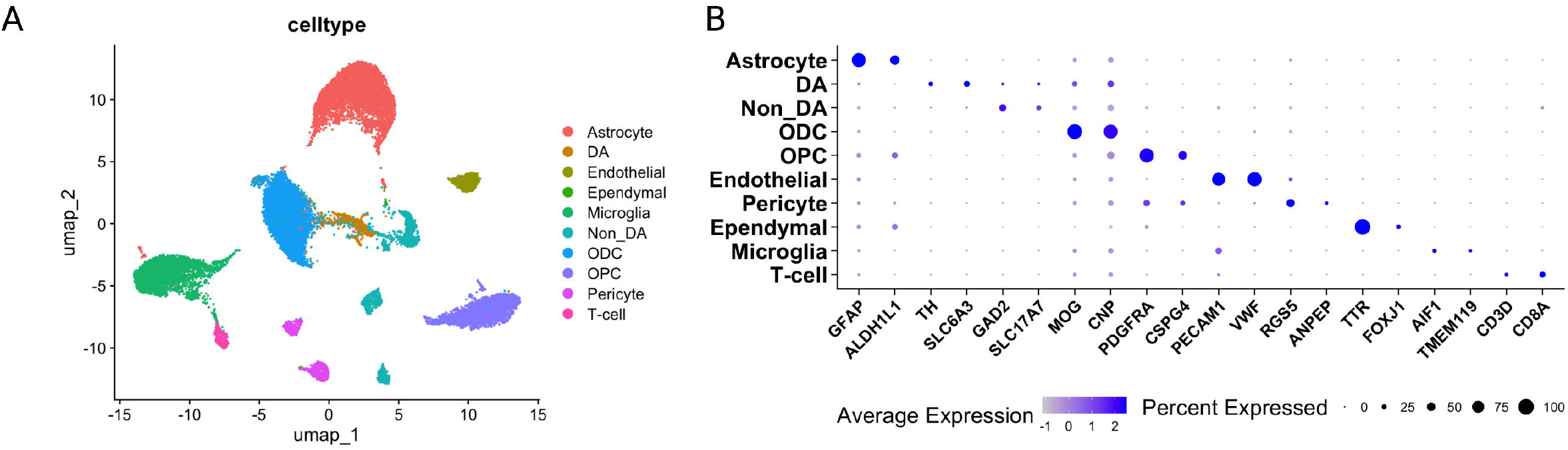

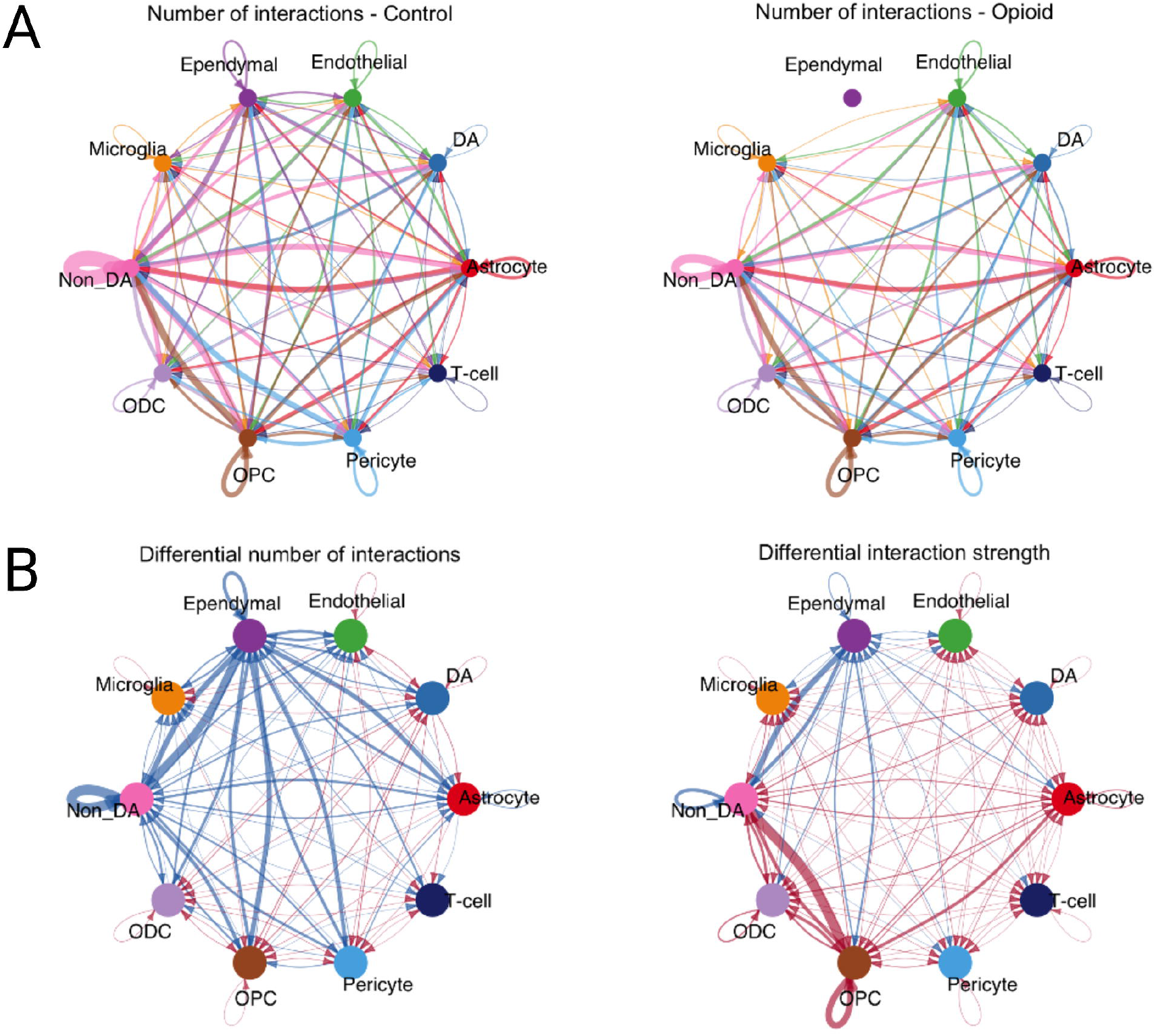

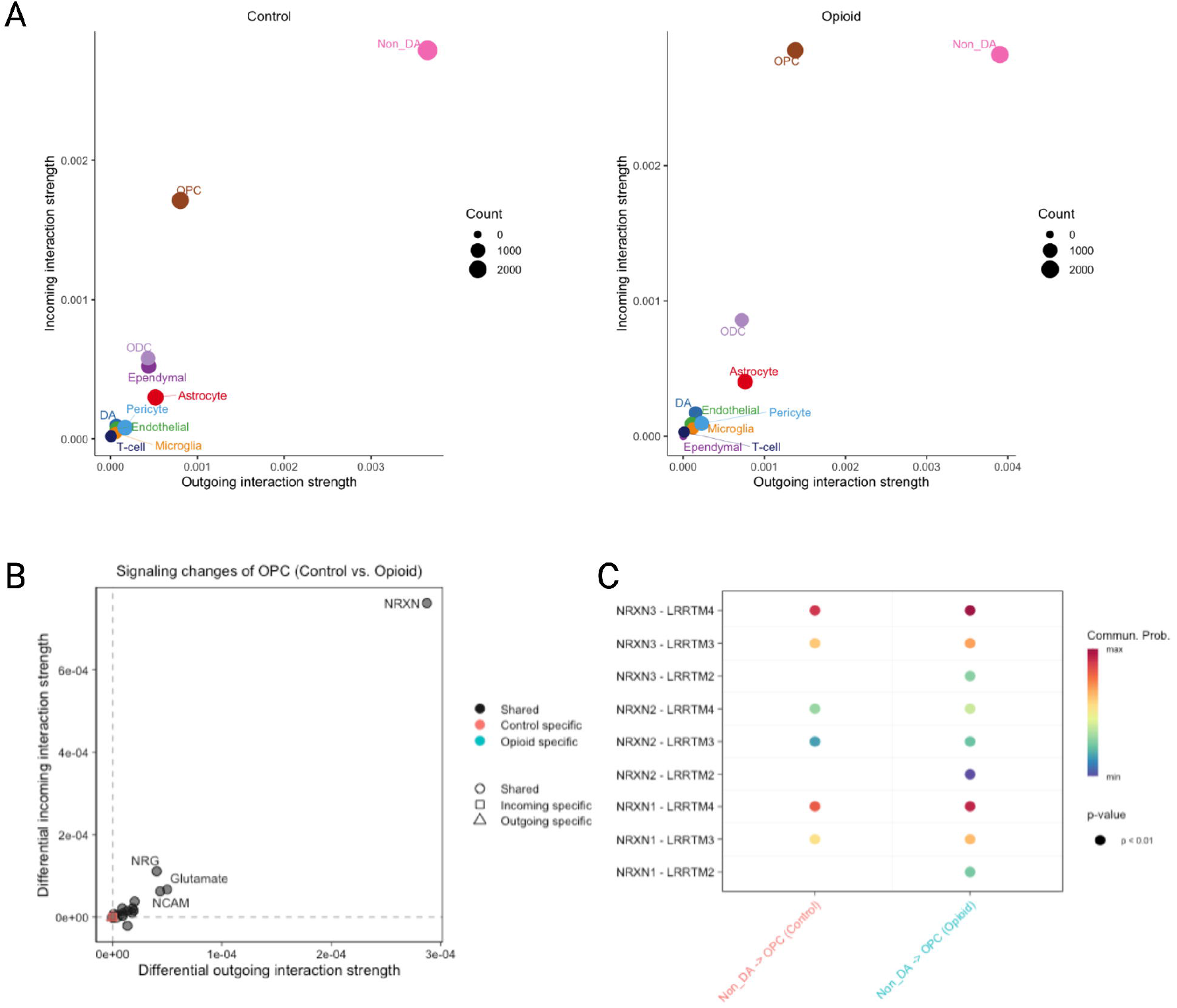

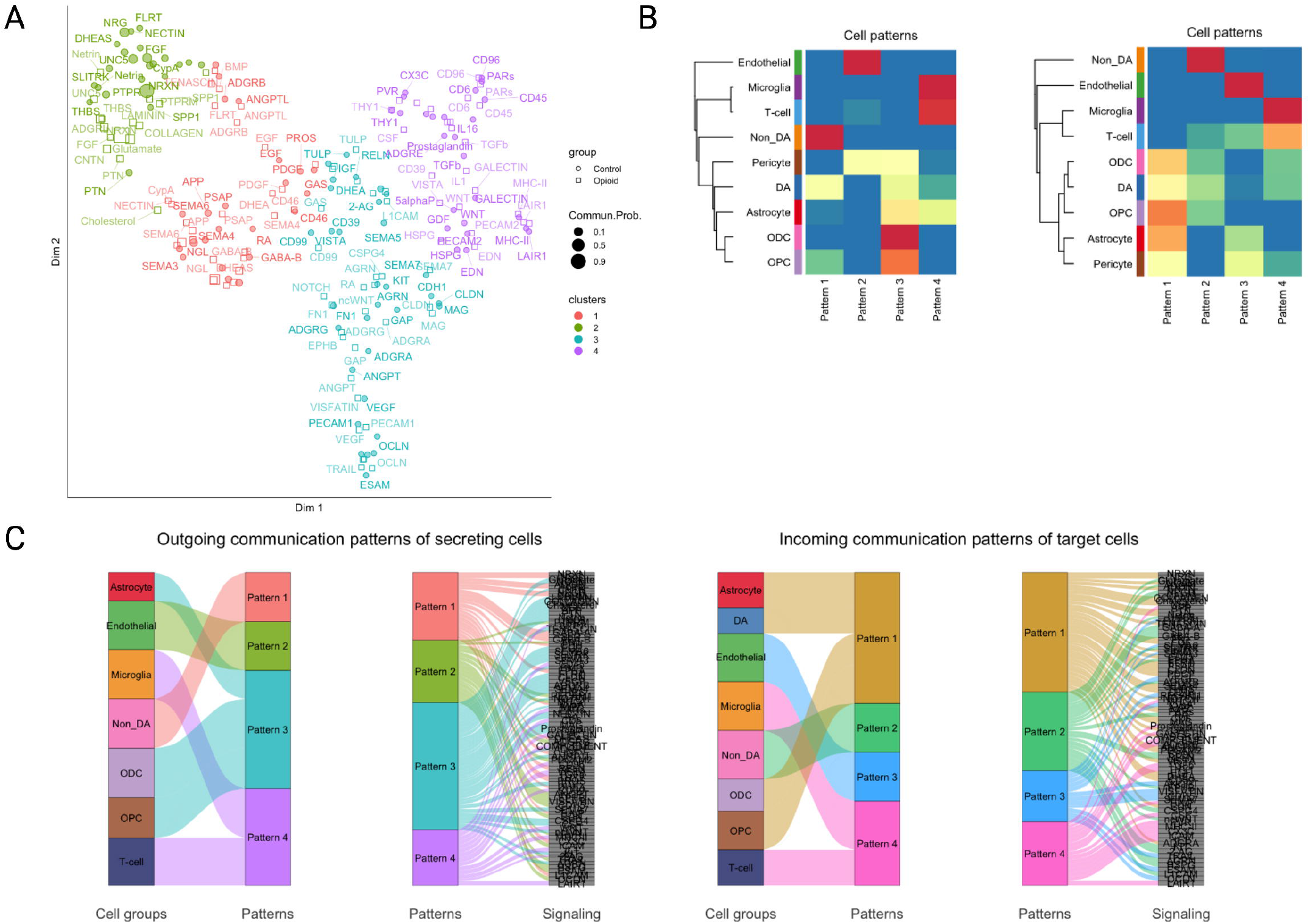

